# Sex-specific declines in cholinergic-targeting tRNA fragments in the nucleus accumbens in Alzheimer’s disease

**DOI:** 10.1101/2023.02.08.527612

**Authors:** Dana Shulman, Serafima Dubnov, Tamara Zorbaz, Nimrod Madrer, Iddo Paldor, David A. Bennett, Sudha Seshadri, Elliott J. Mufson, David S. Greenberg, Yonatan Loewenstein, Hermona Soreq

**Affiliations:** The Edmond & Lily Safra Center for Brain Sciences, The Hebrew University of Jerusalem, Jerusalem 9190401, Israel; The Institute of Life Sciences, The Hebrew University of Jerusalem, Jerusalem 9190401, Israel; The Rachel and Selim Benin School of Computer Science and Engineering, The Hebrew University of Jerusalem, Jerusalem 9190401, Israel; The Neurosurgery Department, Shaare Zedek Medical Center, Jerusalem 9103102, Israel; Rush Alzheimer’s Disease Center, Rush University Medical Center, 600 South Paulina, Suite 1028, Chicago, IL 60612, USA; UT Health Medical Arts & Research Center, San Antonio, TX 78229, USA; Barrow Neurological Institute, St. Joseph’s Medical Center, Phoenix, AZ, 85013, USA; The Department of Neurobiology, The Hebrew University of Jerusalem, Jerusalem 9190401, Israel; The Department of Cognitive Sciences, The Hebrew University of Jerusalem, Jerusalem 9190401, Israel; The Federmann Center for the Study of Rationality, Jerusalem 9190401, Israel

## Abstract

**Introduction:** Females with Alzheimer’s disease (AD) suffer accelerated dementia and loss of cholinergic neurons compared to males, but the underlying mechanisms are unknown. Seeking causal contributors to both these phenomena, we pursued changes in tRNA fragments (tRFs) targeting cholinergic transcripts (CholinotRFs).

**Methods:** We analyzed small RNA-sequencing data from the *nucleus accumbens* (NAc) brain region which is enriched in cholinergic neurons, compared to hypothalamic or cortical tissues from AD brains; and explored small RNA expression in neuronal cell lines undergoing cholinergic differentiation.

**Results:** NAc CholinotRFs of mitochondrial genome origin showed reduced levels that correlated with elevations in their predicted cholinergic-associated mRNA targets. Single cell RNA seq from AD temporal cortices showed altered sex-specific levels of cholinergic transcripts in diverse cell types; inversely, human-originated neuroblastoma cells under cholinergic differentiation presented sex-specific CholinotRF elevations.

**Discussion:** Our findings support CholinotRFs contributions to cholinergic regulation, predicting their involvement in AD sex-specific cholinergic loss and dementia.

## Introduction

AD is the most common cause of dementia, globally affecting over 50 million people^1^. Its clinical symptoms reflect accelerating failure of regulatory, neurochemical and neural networks, initiating with the cholinergic system and leading to a progressive cognitive deterioration. Neuropathologic features in subcortical nuclei^2^ precede neurofibrillary tangles (NFTs) development within the mesial temporal cortex^3^. AD lesions involve accumulation of multimeric amyloid-β fibrils forming neuritic plaques and NFTs consisting of hyperphosphorylated tau and leading to loss of synapses, dendrites, and eventually, neurons^1^. However, the mechanisms underlying AD-related dementia, which develops relatively late in the disease course^4^ are a matter of controversy^5^.

Clustered cholinergic neurons in the *substantia innominate*/nucleus basalis of Meynert^6^ are the most investigated site of acetylcholine(ACh) producing interneurons within the nucleus accumbens (NAc). They contribute to cognitive and motivational behavior and loss of these neurons predicts progression from mild cognitive impairment (MCI) to AD. This is faster in females even when considering their longer life expectancy^7–9^. Females score lower than males on tests of cognitive domains^10^, even after controlling for possible demographic and genetic factors, and show more pronounced mitochondrial imbalances^11^ and loss of cholinergic neurons^12^.

The reduced cholinergic activity in basal cholinergic forebrain neurons and their cortical projection sites supports the ‘cholinergic hypothesis of AD’^13^, which argues that dysfunction of ACh-producing brain neurons contributes to the AD-related cognitive decline. Consequently, the development of cholinesterase inhibitors (ChEI) is one of the few treatments for AD^14^. ChEIs do not prevent or slow disease progression. They may independently lead to cognitive dysfunction and delirium^15^, are more effective in males and cause more pronounced adverse effects in females with AD^7,16^. Additionally, muscarinic agonists have shown some clinical promise, and modulate various impairments seen in AD, affecting both the cortical cholinergic basal forebrain system and the 1–2% of NAc ACh-producing neurons^17^ that project to brain regions including cortical areas^18^. Correspondingly, dysfunction of cholinergic pathways in both the basal forebrain and the NAc contribute to cognitive decline in AD^19,20^.

Both whole tissue and single cell AD studies revealed changes in coding and non-coding RNA (ncRNA) transcript levels, but their involvement in cholinergic cellular dysfunction remained unknown. Those included microRNAs (miRs), small single-stranded ncRNAs about 22 nucleotides long that target mRNA molecules and function as post-transcriptional regulators^21^. Therefore, individual miR levels are co-altered in the AD brain with their experimentally validated targets^22^. Furthermore, sex-specific miR profiles modulate responses to tau pathology^23^ and sex- and age-related ACh signals^24^ in AD, yielding sex-specific biomarkers for multiple neurodegenerative disorders^25^. However, nucleus accumbens AD miRs remain under-investigated.

Transfer RNA (tRNA)-derived fragments (tRFs) are a recently rediscovered small non-coding RNA regulators that were previously considered to be inactive products of tRNA degradation. TRFs are derived by specific tRNA cleavage of several different nucleases, including Dicer, Drosha and Angiogenin^26^ (Figure 1). TRFs up to 30 nucleotides in length may function like miRs by modulating the expression of mRNA targets carrying complementary sequence motifs^26^. In nucleated blood cells, tRFs targeting cholinergic transcripts replace miRs in regulating cholinergic reactions during ischemic stroke^27^. Moreover, certain tRFs regulate ribosomal function, cancer, innate immunity, stress responses, and neurological disorders^28^. TRFs may originate from nuclear or mitochondrial genomes^26^ and mitochondrial disfunction is a feature of several neurodegenerative diseases, including AD^8^. While AD involves changes in miRs targeting mitochondrial genes^29^, the role of mitochondrial-originated tRFs in cholinergic neurons, as well as their sex-specific profiles and genetic origins in AD have not yet been investigated.

**Figure 1:**
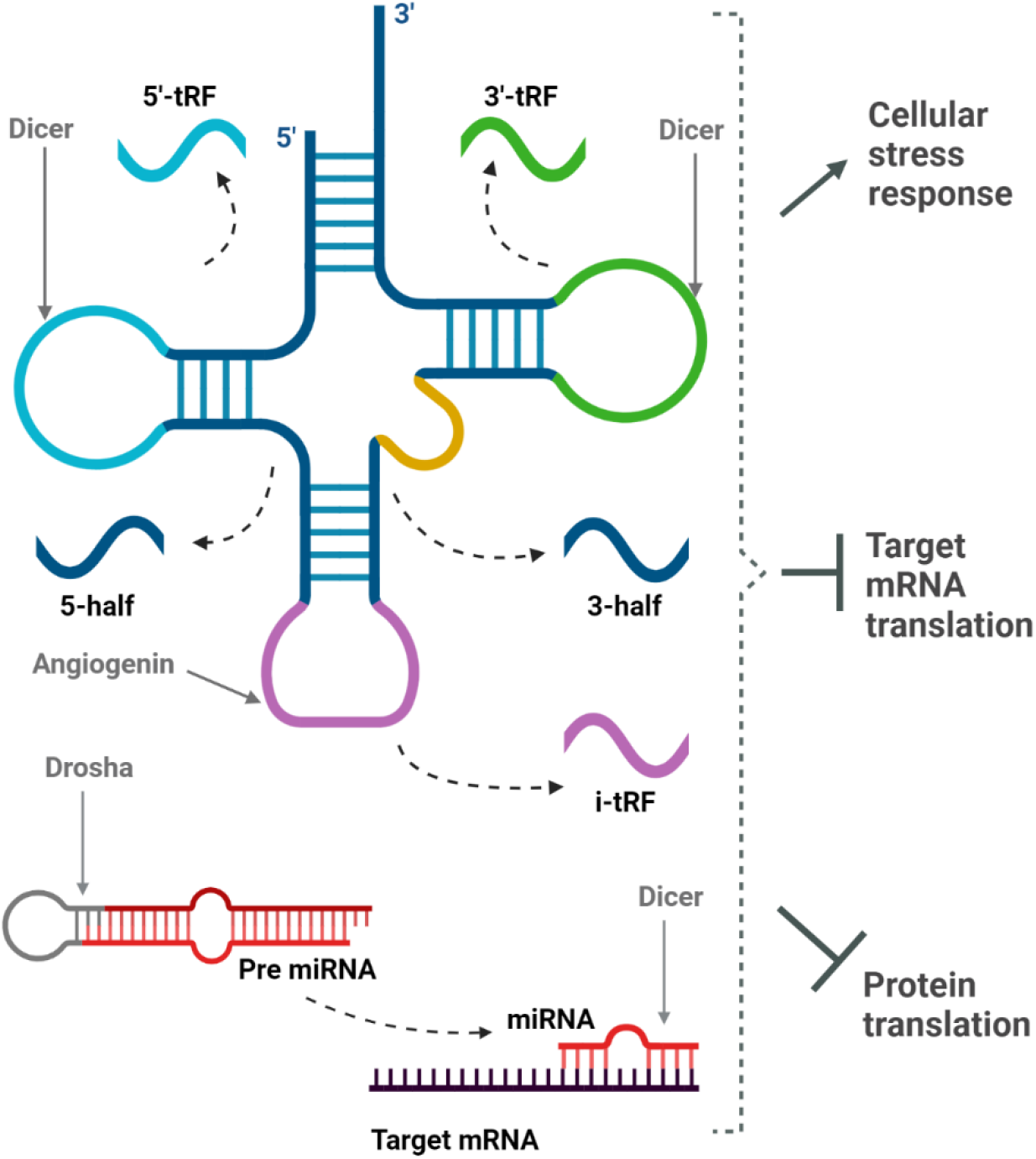
Selected mechanisms in miRs and tRFs.

Pursuing cholinergic-targeting tRFs and/or miRs (CholinotRFs, CholinomiRs) that may affect AD pathology and its disease course and cognitive decline, we sought changes in their predicted mRNA targets in AD and healthy non-cognitively impaired elderly control cases; examined the specific cell populations that express these cholinergic mRNA targets in single cell RNA-Seq from the middle temporal gyrus (MTG), and studied the role of CholinotRFs in cholinergic regulation in cell lines of human origin.

## Methods

### Cohorts

#### RADC Subjects

RNA-Sequencing (RNA-Seq) data derived from participants in the Rush Alzheimer’s Disease Center (RADC) ROS and MAP cohorts^30^. Both studies were approved by an Institutional Review Board of Rush University Medical Center and all participants signed informed and repository consents and an Anatomic Gift Act. All participants in ROSMAP enroll without dementia and agree to annual clinical evaluation and brain donation at the time of death. All cases received a neuropathological evaluation based upon Braak staging, Cerad and NIA-Reagan criteria as well as a semi-quantitative analysis of the number of neurofibrillary tangles (NFT) and neuritic plaque. Post-mortem brain tissues were collected from the noted brain regions of over 65 years old adults without known dementia at enrollment, recruited for the ROS and MAP projects; 196 post-mortem samples were from the NAc (119 females, 77 males), 71 from the STG (36 females, 35 males) and 181 from the hypothalamus (120 females, 61 males).

The condition of the participants was set separately for each trait; cognitive state was set by the clinical diagnosis at time of death (NCI: cogdx score of 1; MCI: 2; AD: 4), neurofibrillary tangle (NFT) state by the semiquantitative measure of severity of NFT pathology (NCI: Braak score of 0-2; MCI: 3-4; AD: 5-6) and the neuritic plaque state was set by the semi-quantitative measure of neuritic plaques (NCI: Cerad score of 1-2; AD: 3-4; in accordance with the recommendation for binary division)^31–33^ (Table S1). Libraries constructed from total RNA were subjected to small RNA-Seq (NEBNext Multiplex Small RNA library prep set for Illumina, New England Biolabs), and sequenced using the Illumina NextSeq 500 platform. Presequencing Bioanalyzer 6000 Quality control was performed using FastQC, and served for adapter trimming and quality-based filtering of all raw reads, and to determine RNA Integrity Number (RIN) tissue values. Samples with RIN < 5 were filtered out. Small RNA was aligned to miRBase using miRexpress and to the tRNA transcriptome using the MINTmap. Long RNA was aligned to the human reference transcriptome for the NAc and hypothalamic samples, but not the STG (ENSEMBL GRCh38 release 79).

#### STG Long RNA-Sequencing

Long-RNA STG sequencing files were obtained via controlled access to the AD Knowledge Portal (https://www.synapse.org/#!Synapse:syn2580853/files/). These data were generated from postmortem STG of 377 volunteers (242 females, 135 males) provided by Dr. Eric Schadt from Mount Sinai School of Medicine, NY. Samples with RIN < 5 were filtered out, leaving 211 samples (136 females, 75 males). Clinical dementia rating scale (CDR) was used to classify subjects as no cognitive deficits (CDR=0) and AD (CDR=3-5), leaving 155 samples (98 females, 57 males).

#### scRNA-Seq data

Cellular level transcriptomic data from the middle temporal gyrus (MTG) of female and male aged volunteers on the AD spectrum were downloaded from the Seattle Alzheimer’s Disease Brain Cell Atlas (SEA-AD) and was accessed in December 2022 from https://registry.opendata.aws/allen-sea-ad-atlas. Study data were generated from postmortem brain tissue obtained from the MTG of 84 aged individuals spanning the full spectrum of AD severity (preMCI, MCI, mild-moderate severe AD) and 5 neurotypical aged adult cognitively intact individuals.

### Data Preprocessing

Raw counts were normalized using the Deseq2’s median of ratios method in which counts are divided by sample-specific size factors determined by median ratio of gene counts relative to geometric mean per gene^34^. The median expression threshold for tRFs and miRs from each brain region was calculated separately for females and males and set as the number of significant features fixed in the maximal number of analyses; 9 (Table S17).

### Statistical Analysis

Kolmogorov–Smirnov test was used to calculate the P-value of each small RNA (miR/tRF) in each tissue and group independently using the SciPy.stats.kstest python function. Benjamini-Hochberg test for false discovery rate (FDR) was used to correct for multiple comparisons (*α* = 0.1) using the statsmodels.stats.multitest python library. The binomial probability was calculated using SciPy.stats. binomtest python function, with alternative=‘greater’. Fisher’s Exact P-values were calculated by SciPy.stats.Fisher_exact.

### Target prediction

The predicted targets of the identified tRFs were determined using the prediction algorithm of the MR-microT DIANA tool based on sequence motif^35^. The likelihood of binding the 3’-untranslated region (UTR) and protein coding sequences (CDS) of each targeted mRNA was calculated separately, combined to a prediction score that was normalized by the conservation of the binding site in a number of species. Targets with prediction score < 0.8 were extracted. The predicted targets of the miRs were found using the DIANA-microT-CDS prediction algorithm^35,36^ to enable the best comparison between tRF and miR targets. Predicted targets with a prediction score < 0.8 were extracted. The python libraries selenium and multiprocessing were used for web scraping in order to automate the use of the DIANA prediction tools.

### Definition of cholinergic sncRNAs

Literature review enabled compilation of 58 cholinergic system related genes divided into two main pathways^37–50^; ACh-production and cholinergic regulation (Table S4). The cholinergic threshold for sncRNAs was set at the 80^th^ quantile of the number of ACh production or cholinergic regulation-associated targets in each sample independently, for females and males combined (tRFs; 4, miRs; 5, for the NAc, STG and hypothalamus), based on the work in Lobentanzer et al., 2019^43^. sncRNAs passing the cholinergic threshold, or targeting at least one ‘core’ cholinergic gene (Table S4, denoted as ‘Core’), were defined as CholinotRFs / CholinomiRs.

### Logistic Regression for interaction analysis

The Logit model from statsmodels.discrete.discrete_model was used for interaction analysis between two discrete variables; the potential function of the tRF (CholinotRF vs. non-CholinotRF, denoted by 1 and 0, respectively) and the origin of the tRF (MT vs. non-MT, denoted by 1 and 0, respectively). The label of each tRF represented the direction of change in AD state (Reduction vs. Elevation, denoted by 1 and 0, respectively). The coefficients and their significance represented the correlation of these variables to the change in AD state.

### scRNA-Seq analysis

We analyzed the scRNA-Seq dataset from the Seattle Alzheimer’s Disease Brain Cell Atlas (SEA-AD) study. Preprocessing and analysis of snRNA-Seq data was implemented with the SCANPY package, on the entire cohort, including 1,378,211 cells and 36,601 genes. Cholinergic genes with median expression smaller than 100 in the STG bulk RNA-Seq data were excluded, leaving 31 cholinergic genes (Table S4).

Linear regression was applied to STG bulk RNA-Seq data and the linear combinations representing the expression of each gene as the sum of the expression in each cell type population and the relative number of each cell population from the entire cohort (the numbers of the cells are listed in Table S10). Seaborn.regplot was used for the linear regression plot and SciPy.stats.linregress was used for significance calculation with alternative=‘greater’. Expression values of the cells were calculated according to the scanpy.tl.rank_genes_groups method: np.log2((np.expm1(AD mean) + 1e-9) / (np.expm1(Control mean) + 1e-9)). The log fold changes of each gene in each cell population can be found in Tables S11-S12, in which the significance of the change was calculated using scanpy.tl.rank_genes_groups with method=‘wilcoxon’. The correlation coefficients were calculated using SciPy.stats.stats.pearsonr.

### Cell culture

The Cell Lines LA-N-2 (female) (DSMZ Cat# ACC-671, RRID:CVCL_1829) and LA-N-5 (male) (DSMZ Cat# ACC-673, RRID:CVCL_0389) were purchased at DSMZ (Braunschweig, Germany). These cells respond to differentiation by several neurokines (CNTF, LIF, IL-6) by cholinergic differentiation according to a known protocol^51^, corresponding with elevation of choline acetyltransferase (ChAT), the central cholinergic marker (mRNA, protein, and activity) as well as it’s intronic vesicular ACh-transporter gene, vAChT (aka SLC18A3). The small-RNA sequencing files were produced in-house and are available to all interested researchers from the GEO data portal (https://www.ncbi.nlm.nih.gov/geo/query/acc.cgi?acc=GSE132951, GEO accession: GSE13295). Each time point (2 days vs 4 days) contained 4 replicates from each state (Control vs Differentiated) for each sex (Female vs Male).

CholinotRFs were defined in the same way as they were defined for the brain samples (cholinergic threshold: 3). SciPy.stats.ttest_ind was used for the calculation of the t-test P-value for the presentation of the tRF changes in cell lines. The results are reported in Tables S13-S16.

## Data and code availability

The ‘Sex_specific_declines_2023’ repository holds the data and code information: https://github.com/danashulman/Sex_specific_declines_2023.git.

## Results

### NAc CholinotRFs decline in AD females

To identify disease- and sex-specific changes in the NAc sncRNA profiles, we analyzed small RNA-Sequencing (RNA-Seq) data of aged donors with varying degrees of cognitive impairment. The cholinergic hypothesis of AD states that dysfunction of ACh-producing brain neurons contributes to the cognitive decline. Therefore, we classified the donors according to their clinical diagnosis: controls with no cognitive impairment (NCI) (65; 35 females, 30 males) and persons diagnosed with AD (47; 28 females and 19 males) (*Methods*, Table S1)^31–33^. To find AD-associated changes in short tRFs, we applied a Kolmogorov–Smirnov test on control and AD short RNA-Seq reads (*Methods*). We found substantial differences in the levels of sncRNAs between males and females. Specifically, in females, we identified 10 tRFs with altered levels (FDR corrected P≤0.1^52^) (Figure S1, Table S2). In males, by contrast, no significant alterations in tRFs were found (Figure 2A).

**Figure 2:**
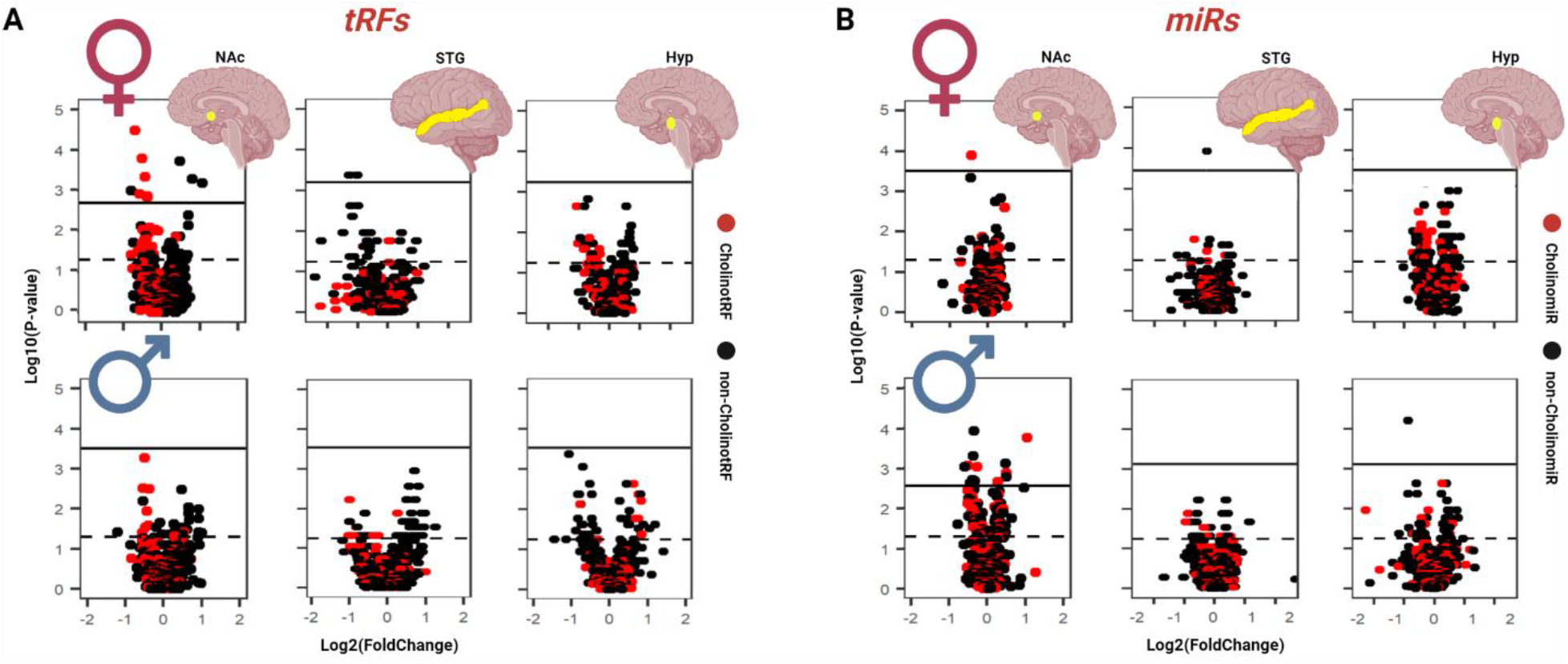
CholinotRF reductions were most significant in the NAc of AD females. Volcano plots presenting changes of the levels of cholinergic sncRNAs the predicted NAc, STG and Hypothalamus mRNA targets of whom contribute to ACh production and cholinergic regulation in cognitively impaired group compared to controls. Red: sncRNAs with predicted cholinergic activities. Black: other sncRNAs. Solid line: the significance threshold of the FDR corrected P-value. Dashed line: Kolmogorov Smirnov test P of 0.05. A) Changes in CholinotRFs. B) Changed in CholinomiRs. Top row: Changes in females. Bottom row: Changes in males.

AD involves the accumulation of neuro-fibrillary tangles (NFT) and neuritic plaques. However, their link to cognitive decline is unclear^4^. Therefore, we asked to what extent changes in short tRFs and miRs will manifest when AD is classified according to these neuropathological features, rather than the level of cognitive decline (Table S1). When classifying the donors according to neuritic plaques we found no significant differences in the levels of sncRNAs (Figure S2A, *Methods*). When classifying them according to NFTs, we identified only a single tRF that significantly differed between the two groups (Figure S2B, Table S3, *Methods*). These results support the use of cognitive impairment rather than neuropathology as a measure of disease severity.

Based on the cholinergic hypothesis, we hypothesized that the AD-related changes in tRFs will be particularly pronounced in those tRFs containing complementary sequences to cholinergic genes. To test this hypothesis, we assembled a list of 58 genes associated with ACh production and cholinergic regulation (Table S4, denoted as ‘ACh Production’ or ‘Cholinergic Regulation’^37–50^, based on our previous work^43^), of which 7 genes were denoted as ‘Core’^38,43^ genes based on their key roles in the cholinergic pathway (Table S4). For each sncRNA, we used the microT prediction tool to find its cholinergic targets^35^. This identified approximately a quarter of sncRNAs likely to be involved in ACh production and cholinergic regulation (*Methods*). Focusing on tRFs whose level was significantly altered in AD compared to control state, 6 of the 10 tRFs altered in females were CholinotRFs, while only 25% of all the tRFs (98/401) were identified as CholinotRFs (P<0.02) (*Methods*).

The levels of all 6 significant NAc CholinotRFs were reduced in AD females (P<0.02). We were therefore wondering whether this reduction is only the tip of an iceberg and reflects a more general reduction in the level of CholinotRFs, including in those tRFs that did not cross the significance level. Indeed, in females, the level of 74% (68/92) of the remaining (no significant change) CholinotRFs was lower in the AD than in the control state, while the level of only 26% (24/92) was higher (P<3×10^−6^). By contrast, no such reduction was observed in the non-CholinotRFs (36%, 108/303 decreased; 64%, 195/303 increased). A similar tendency, to lesser extent, was also observed in males: 70% (68/97) of CholinotRFs decreased and 30% (31/97) of CholinotRFs increased (P<5×10^−5^); 47% (135/288) of non-CholinotRFs decreased and 53% (153/288) of non-CholinotRFs increased.

Caution should be exercised when interpreting this result, because the classification of tRFs into CholinotRFs and non-CholinotRFs is correlated with their origin, mitochondrial (MT) or nuclear (non-MT). For example, in females, 82% (80/97) of the CholinotRFs are also MT-tRFs, whereas only 10% (29/303) of the non-CholinotRFs are also MT-tRFs. We therefore used logistic regression to test whether the potential function of a tRF (CholinotRF vs. non-CholinotRF) is correlated with its reduction in AD, even when its origin (MT vs. non-MT) is accounted for (*Methods*). The regression coefficients indicate that both in females and in males, the potential function of a studied tRF as cholinergic regulator is correlated with its reduction in the AD NAc (Females: regression coefficient 1.1, P<1×10^−4^; Males: regression coefficient 1.1, P<9×10^−4^), whereas the regression coefficient for the origin was not statistically significant (Females: regression coefficient 0.006, P<0.98; Males: regression coefficient −0.25, P<0.35), indicating that the effect in MT-tRFs derived from their potential function as cholinergic regulators.

### tRFs decline is more pronounced in the NAc than in other brain regions

It is important to note that only a small fraction of NAc neurons is cholinergic, limiting the interpretation of the RNA-Seq data that was derived from a heterogenous population of NAc neurons. To test whether the change in tRFs reflects changes in the NAc cholinergic neurons, we repeated the analysis in two additional brain regions, the STG and hypothalamus. The STG is a cortical region that was previously linked to AD, presenting epigenetic modifications in persons with AD according to their pathogenesis^53^, whereas the hypothalamus is a subcortical region that is relatively spared^54^. We classified small-RNA profiles extracted from the STG and hypothalamus of controls (STG: 11 females, 11 males; Hypothalamus: 15 females, 11 males) and donors diagnosed with AD (STG: 10 females, 6 males; Hypothalamus: 20 females, 10 males) (Table S1). In the STG, we identified only 2 tRFs whose levels are significantly changed in females between the control and AD groups. None of these tRFs was a CholinotRF (Table S5). In the hypothalamus, we did not find any tRFs that differed between the two groups. In agreement with these results, the reduction in the global level of CholinotRFs was modest, statistically significant only for males in the STG and females in the hypothalamus (STG Females: 57%, 55/97, P<0.11, Males: 62%, 65/104, P<7×10^−3^; Hypothalamus Females: 58%, 54/93, P<0.07; Males: 40%, 36/92, P<0.99). These findings lend support to the potential cholinergic relevance of our NAc observations. While the dorsomedial portion of the hypothalamus contains cholinergic neurons^54^, our findings in the hypothalamus part that is non-cholinergic were compatible with the link between the altered tRFs in the NAc and their functional relevance to cholinergic neurons.

### AD-related changes in miR levels differ substantially from AD-related changes in tRFs

Many studies have reported post-transcriptional regulation of gene expression by miRs^14,21,27^. Therefore, if the changes in tRFs merely reflect changes in the fraction of cholinergic neurons (as a result of cell death) then a similar pattern of change in miRs is expected in AD. However, miRs exhibited very different behaviors from those of tRFs. While substantial changes in the levels of tRFs were observed in the in NAc of AD females, but not males, the opposite pattern was observed in miRs. In males we identified 13 miRs whose levels were significantly altered between the AD and control groups (Figure 2B, Table S6), while only a single miR was significantly altered in females (Table S7). Intriguingly, the single significantly reduced miR in the NAc of AD females was a CholinomiR (the well-studied AChE-targeting miR-132-3p^24^), while 5 of the 13 significantly altered miRs in AD males were CholinomiRs. Similarly inverse changes were seen in the population of CholinomiRs (*Methods*): while 84 of 144 (58%) CholinomiRs were reduced in the NAc of AD males (P<0.03), 56% of CholinomiRs in the NAc of females were elevated (80/143, P<0.09).

In the STG, we identified only a single miR, which was not a CholinomiR whose level was significantly reduced in AD females (Figure 2B, Table S8). At the population level, 58% (93/160) of the CholinomiRs were significantly reduced in AD females (P<0.03), while CholinomiRs reduction was not significant in AD males (50%, 84/168, P<0.53). In the hypothalamus, we identified a single miR, which was not a CholinomiR, whose level was significantly reduced in AD males (Table S9). At the population level, CholinomiRs were more significantly reduced in females (62%, 85/138, P<4×10^−3^) than in males (56%, 79/141, P<0.09).

Taken together, these findings point at a complex network of gene expression regulation that differs between tRFs and miRs, between females and males and between brain regions. We find substantial sex-specific differences in the regulation of the cholinergic system in the NAc. These include substantial changes in CholinotRFs in AD females, but almost no changes in CholinomiRs, while the opposite trend, substantial changes in miRs and less so in CholinotRFs, were observed in males with AD.

### Low CholinotRF levels in the NAc of AD females are correlated with elevated levels of cholinergic mRNAs

Short tRFs are likely to interact with and suppress the translatability of mRNAs carrying complementary sequences^21,22,26^ (Figure 1). Therefore, we hypothesized that a reduction in the levels of CholinotRFs in the NAc of AD females will be accompanied by elevated levels of cholinergic mRNAs. To test this prediction, we used a permutation test to identify those genes, out of the 58 genes associated with ACh production and cholinergic regulation^37–50^ (Table S4), whose levels were significantly modified in AD compared to control groups (*Methods*). Consistent with our hypothesis, the levels of 93% (13/14 genes) of the significantly altered mRNAs associated with ACh-production and cholinergic regulation were elevated in females (P<9×10^−4^). In comparison, in males only 7 genes displayed altered levels, and they were all elevated (P<8×10^−3^).

Next, we focused our attention on the specific CholinotRFs that exhibited a significant reduction in the NAc of AD females (6/98). Of these 6 tRFs, 5 shared similar targets. Therefore, while each of these CholinotRFs targeted 15-16 cholinergic targets, together they targeted only 18 targets (out of the 58 cholinergic genes). This similarity in targets was due to the fact that all 5 tRFs originated from the same tRNA (mitochondria-derived Phe^GAA^) (Figure 3A, Table S2, *Methods*). We hypothesized that the levels of these 18 targets will be particularly affected in females with AD. Indeed, the mRNA levels of 33.3% (6/18) of the cholinergic targets of these CholinotRFs were significantly increased in AD females compared to controls. Considering the remaining 40 cholinergic genes, only 17.5% (7/40) exhibited a significant increase in AD females (P<0.08). In comparison, in males only 6% (1/18) of the target genes and 15% (6/40) of the non-targeted genes were significantly increased.

**Figure 3:**
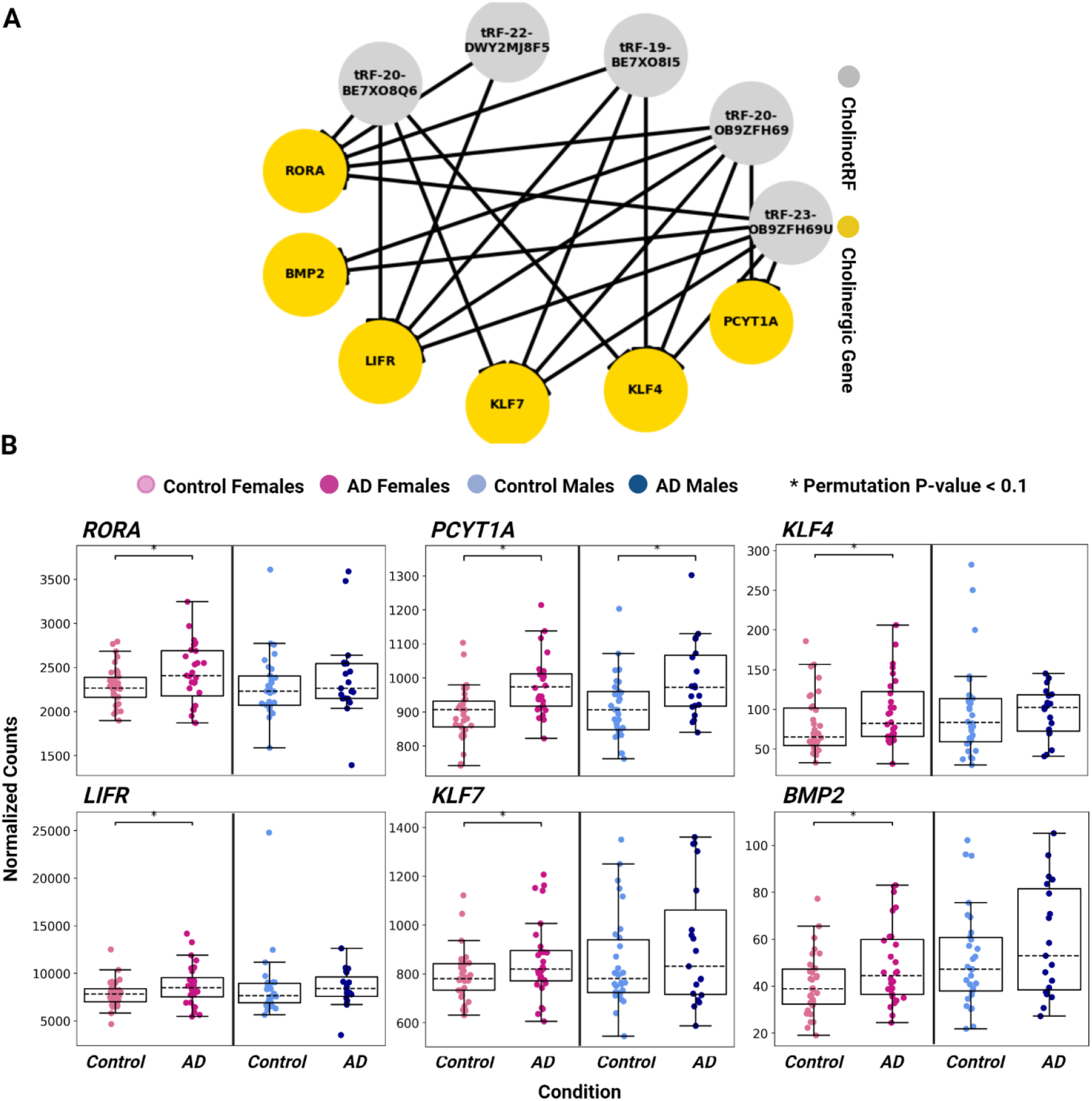
Low CholinotRF levels in the NAc of AD females are correlated with elevated levels of cholinergic mRNA targets associated with ACh production and cholinergic regulation. A) Network of potential regulation links in female brains. Grey nodes: decreased CholinotRFs of interest in AD females. Yellow nodes: elevated cholinergic transcripts in AD females. Edge: sequence-based predicted targeting of the cholinergic transcript by the CholinotRF. B) Normalized levels of the cholinergic target mRNAs of tested CholinotRFs. Pale pink: Female controls. Dark pink: AD females. Pale blue: Male controls. Dark blue: AD males. * Changes with Permutation P-value < 0.1.

Previous studies have linked the 6 significantly elevated target mRNAs reported here with AD or related pathways. For example, RORA is elevated in the AD hippocampus^55^, and is genetically linked to AD^56^ (P<0.06, 0.39 for females and males, respectively); PCYT1A is elevated in the AD hippocampus compared to controls^57^ (P<3×10^−4^, 0.01); KLF4 and LIFR were suggested to serve as AD therapy targets based on bioinformatic analyses of therapeutic pathways^58,59^ (KLF4: P<0.09, 0.90; LIFR: P<0.06, 0.98); also, RORA, KLF4 and KLF7 (P<0.06, 0.27) are regulators of the circadian rhythm^42,60,61^, a process that affects cholinergic differentiation and activity^19^, and which notably deteriorates in AD; and BMP2 regulates the Wnt/*β*-catenin pathway which is dysfunctional in several diseases, including AD^62^ (P<0.03, 0.22). Therefore, these findings supported our working hypothesis that sncRNAs in the NAc may operate as regulators of cholinergic associated genes and contribute to the female-specific disease expression.

### Unexpected changes in STG and hypothalamus cholinergic mRNAs

Compared with the NAc, AD-associated changes in cholinergic sncRNAs in the hypothalamus and STG were modest. Therefore, we expected to see fewer AD-related changes in hypothalamus and STG cholinergic mRNAs in general and no elevation in these genes in particular. These predictions were only partially confirmed as we did find a substantial number of cholinergic genes whose level significantly differed between the control and AD states. In the hypothalamus, we identified 3 and 8 genes whose levels significantly changed in females and males, respectively (Figure S3). These changes were even more pronounced in the STG, where the levels of 22 and 18 cholinergic genes changed significantly in females and males, respectively (Figure S4). However, consistent with our prediction, there was no significant elevation of cholinergic mRNAs (Hypothalamus: Females: 1/3, P<0.88; Males: 5/8, P<0.37); STG: Females: 14/22, P<0.14, Males: 10/18, P<0.41).

### Altered levels of STG cholinergic transcripts can be explained by changes in distinct cell populations

The altered levels of cholinergic sncRNAs and mRNAs found in the NAc of AD cases, and AD females in particular, are attributed to cholinergic neurons in the NAc^17^. However, this would not explain the altered levels of cholinergic mRNAs found in the hypothalamus and STG, since these regions lack cholinergic neurons^63^. Nevertheless, previous studies identified AD-related cholinergic changes in other cell populations, including oligodendrocytes^64^, astrocytes^65^ and microglia^14^. Therefore, we performed studies to identify the cell types that contributed to the altered levels of cholinergic mRNAs seen in these extra NAc regions. Specifically, we used single-cell RNA-Sequencing (scRNA-Seq) data from the Seattle Alzheimer’s Disease Brain Cell Atlas (SEA-AD) study (*Methods*), to determine cell type-specific changes in cortical neurons. To pursue the potential cell types where such changes could occur, we compared Medial Temporal Gyrus (MTG, which is located next to the STG in the brain, Figure 4A) data from control and AD brains, as the changes were more pronounced in the STG than in the hypothalamus. The data included classification to 24 distinct cell populations (Figure 4B), of which 18 were of neuronal origin and 6 of non-neuronal origin (astrocytes, oligodendrocytes, OPCs, microglia-PVM, endothelial and VLMC). Notably, the non-neuronal cells constituted only ∼20% (20.7% in females, 19.3% in males) of all the cells in the analysis.

**Figure 4:**
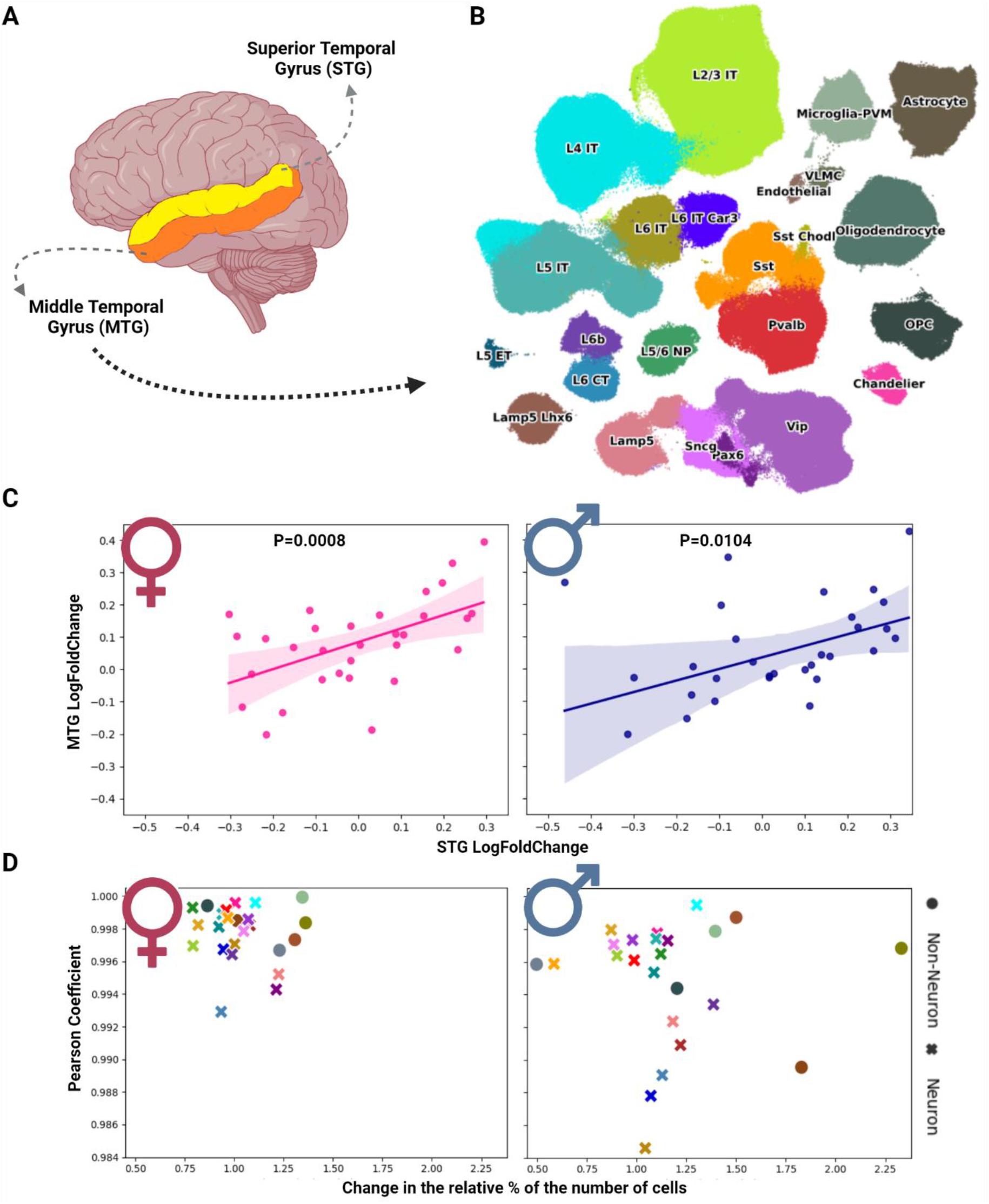
Changes in the levels of mRNAs associated with ACh production and cholinergic regulation are attributed to changes in distinct cell populations. A) The Middle Temporal Gyrus (MTG) in the brain. B) 24 cell populations in the MTG scRNA-Seq data, taken from the SEA-AD study in the Allen Brain Map portal. C) Linear regression applied on the log fold changes of 31 cholinergic genes in STG bulk RNA-Seq and in linear combinations representing the expression in MTG data. Pink: Changes in females. Blue: Changes in males. Dots: 31 cholinergic genes. D) AD-related changes in MTG distinct cell populations. X axis: Change in the relative percentage of the number of cells in AD compared to Control. Y axis: Pearson coefficient on the expression vectors of the 31 cholinergic genes in AD and Control state, separated by sex. Colors: Cell populations as colored in Figure 4B. Dots: non-neuronal cell populations. X symbols: Neurons.

First, we tested whether the MTG scRNA-Seq results are comparable with the STG bulk RNA-Seq. Finding such a consistency is not a trivial result for two reasons. First, while these regions are adjacent, they are not identical and therefore mRNA levels could differ between the two regions. Second, scRNA-Seq measures mRNA levels in the soma, whereas bulk RNA-Seq measures mRNA levels throughout the region, which includes neurites and extensions of non-neuronal cells. We focused on genes that passed the expression threshold (*Methods*) from among the cholinergic genes associated with ACh production and cholinergic regulation (31 genes, Table S4), as previous studies reported unreliable detection of changes in genes with low expression levels in scRNA-Seq^66^. To test for consistency, we represented the expression of each gene in healthy and AD states as a linear combination of the mean gene expression in each cell type and the relative weight of the number of cells in each population (Figure 4C; dots, Table S10). A linear regression on the log fold changes of these transcript levels in the STG bulk RNA-Seq and in the linear combinations representing gene expression in the MTG exhibited a slope that is significantly positive for both females and males (slope=0.42, 0.36; P<8×10^−4^, 0.01 for females and males, respectively) (Figure 4C; solid line, *Methods*), demonstrating that the two measurements are consistent with each other.

With the scRNA-Seq measurements, we can go beyond the bulk RNA-Seq results and study the specific cellular changes associated with the disease. For each population of cells, we computed the change in the *fraction* of cells, a measure of specific cell death (or formation), and the change in the transcript level of cholinergic genes, quantified by the correlation between the vectors of expression in control compared to AD (Figure 4D, *Methods*). In general, the correlation coefficients were relatively high across cell populations, suggesting that the expression levels within a single population were relatively mild. By contrast, there were significant changes in the fractions of cells, indicating vulnerability of specific cell populations within the AD brain. Interestingly, the observed changes were more prominent in males, a result that is consistent with the observation that the cortex is affected more severely in males compared to females in AD^67^.

Of particular interest is the observation of a general increase in the glial cell types (oligodendrocytes, astrocytes and microglia-PVM) in AD females. The opposite trend was observed in males. Notably, while the amounts of glial cells decreased in AD males compared to controls, the decrease in the number of neurons was greater and led to an increase in the relative percentage of glia (Figure 4D, Table S10). For a complete list of changes, see Tables S10-S12.

### CholinotRFs exhibit sex-related elevations during cholinergic differentiation of human-originated neuroblastoma cells

Our definition of CholinotRFs was based on sequence-based complementarity with cholinergic genes. To challenge their role in cholinergic regulation, we studied changes in small RNA-seq transcripts in human neuroblastoma cell lines from male and female origins (LA-N-2, LA-N-5) 2 days after exposure to neurokines that induce cholinergic differentiation^43^ (Figure 5A, *Methods*). We hypothesized that differentiation would lead to elevated levels of all cholinergic RNAs, including CholinotRFs, related to cholinergic production.

**Figure 5:**
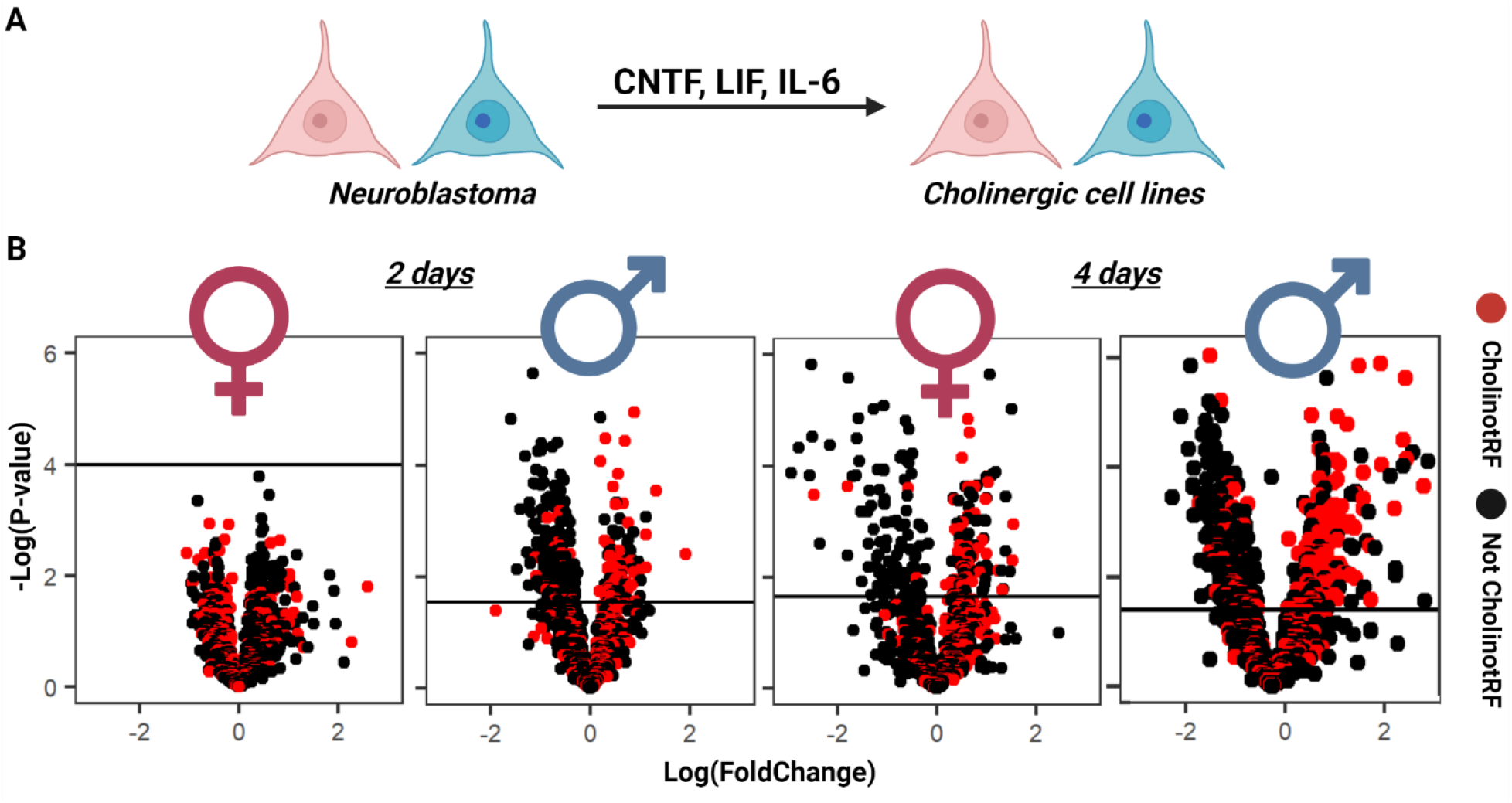
Cholinergic-differentiated cell lines of man and woman origins show elevated levels of CholinotRFs compared to non-differentiated LA-N-2 and LA-N-5 cells. A) Human-originated neuroblastoma cell lines of male and female origins differentiate into cholinergic neurons when exposed to the neurokines CTNF, LIF and IL-6. B) For convenient presentation, t-test was applied on the control and differentiated cell-lines. Volcano plots reflect CholinotRF levels in male and female cholinergic cell lines compared to the original LA-N-2 and LA-N-5 2 and 4 days after differentiation. Red: tRFs with predicted cholinergic activities. Black: other tRFs. Solid line: the significance threshold of the FDR corrected P-value.

Indeed, following 2 days of treatment we found substantial increases in the levels of CholinotRFs in male originated cell lines. Of 308 CholinotRFs passing the expression threshold, the levels of most increased (65%, 201/308, P<5×10^−8^; Table S13). This increase was specific to CholinotRFs, as no such increase was observed in non-cholinergic tRFs (39%, 279/710, Fisher’s Exact P<1×10^−15^; Figure 5B). Surprisingly, we did not observe similar increases in female originated cell lines: the levels of only 38% (118/308) of CholinotRFs were elevated (compared with 35% of non-cholinergic tRFs, 247/710, Fisher’s Exact P<0.29; Figure 5B; Table S14). Since a previous study reported that neuronal cholinergic differentiation is a slower process that matures around 4 days^43^, we further examined tRF levels 4 days after exposure to neurokines. At this time point, CholinotRF levels were elevated in the differentiated cells in both lines (LA-N-2: 77%, 209/273, P<1×10^−10^; LA-N-5: 64%, 174/273 P<3.5×10^−5^; Tables S15-S16). This increase was not found in non-cholinergic tRFs (LA-N-2: 47%, 335/706 non-cholinergic tRFs increased; LA-N-5: 33%, 234/706; Figure 5B). The change was even larger in the female-originated cell line than in the male-originated cell line (P<0.01). Thus, CholinotRFs were lost faster in the NAc of AD females compared to males NAc and were gained more profoundly in female-originated neuroblastoma cells compared to male neuroblastoma cells under cholinergic differentiation.

Taken together, these results strengthen the link between the sequence-defined cholinotRFs and cholinergic regulation. Moreover, they support the finding of sex-related differences in the cholinergic regulation of AD via sncRNAs.

## Discussion

While the cholinergic hypothesis of AD^13^ has received less attention in recent years, the accelerated damage to ACh producing neurons and the cognitive decline in females with AD^19^ call for exploring the molecular processes controlling cholinergic dysfunction. Here, we focused on miRs and tRFs as regulators of ACh production and cholinergic regulation in brain regions and cell types with diverse cholinergic activities. To pursue potential regulators of the more severe and more frequent AD in females compared to males^8^ we compared RNA-Sequencing datasets of NAc, STG and hypothalamus from AD and non-cognitively impaired elderly. We sought sex-specific changes in miRs/tRFs and their target mRNA levels, compared scRNA-Seq patterns derived from a closely related brain region and pursued miR/tRF changes in neuronal cell lines of human male and female origin under cholinergic differentiation.

Greater changes in cholinergic RNAs were found in females with AD compared to males. Overall tRF levels were also reduced in females and the reduction in cholinotRFs correlated with larger increases of cholinergic mRNAs in the NAc of AD females. We also observed changes in cholinergic mRNA levels in non-neuronal cell populations from cortical tissues of AD females; and slower response to cholinergic differentiation that may be causally linked to the exacerbated cognitive decline in AD females^7–9^.

The cholinergic hypothesis posits that loss of cholinergic neurons contributes substantially to the cognitive decline in those with advanced age and AD^13,19^. Here, we found greater AD-related changes in miRs/tRFs regulating cholinergic processes in the female NAc, indicating that small RNA regulators of cholinergic mRNA targets are affected more significantly in the NAc, which is enriched in cholinergic neurons compared to brain regions lacking ACh positive cells such as the STG and hypothalamus^17,63,68,69^. Further, the NAc of AD females but not males revealed region-specific reductions in the levels of tRFs capable of targeting cholinergic genes and an accompanying elevation of cholinergic target transcripts in the NAc (e.g. RORA^42^, BMP2^50^, LIFR^59^, PCYT1A^45^, KLF4^41^ and KLF7^61^), all associated with AD or related pathways. These may temporarily re-balance cholinergic signaling in the NAc of AD females compared to non-cholinergic regions such as the hypothalamus^68^.

The NAc’s role in learning, memory and cholinergic signaling is well established^69^, supporting the pronounced miR/tRF changes in the NAc. Additionally, the observed changes associate well with the cognitive impairment rather than with neurofibrillary tangles and neuritic plaques in both AD females and males, compatible with these regions’ involvement in learning and memory processes and cholinergic signaling^69^. Our findings of modified miR/tRF levels related to the clinical rather than the neuropathological characteristics lends support to the current diagnosis of AD based upon the diagnostic state of the patient reported by clinicians^70^. The null association of cholinergic transcripts with amyloid and tau lesions further suggested a greater role for cholinergic dysfunction in the onset of cognitive decline in AD.

Intriguingly, CholinotRF levels were reduced in the NAc of both AD females and males, albeit to different extents, whereas CholinomiR levels were elevated only in AD females. Inverse reductions of CholinomiRs and CholinotRFs elevations occur in post-stroke nucleated blood cells^27^. However, nucleated blood cells continue to divide, unlike brain neurons, and stroke has a rapid onset and drastic systemic changes, which occur within a short time frame^27^, unlike the slower onset of cognitive decline seen in AD, where rapid production of regulatory miR/tRFs may be unsustainable. Together, our findings suggest that fundamental differences between immunologic responses to rapid brain insults underlie the decreases in CholinotRFs that likely reflect a long-term depletion of the tRF stores in the AD female brain. That we also found AD-linked changes in the NAc levels of miRs in AD males but not females, which potentially indicates an imbalanced miRs regulation in females, highlighting the potential links between miRs and tRFs control and the importance of analyzing disease RNA markers in a sex-specific manner.

Our finding of a reduced levels of MT-tRFs in the AD NAc and their link to cognitive impairments extend previous studies linking mitochondrial impairment to neurodegenerative and psychiatric disorders^8^. Additional studies identified MT-tRFs as key regulators in nuclear-mitochondrial communication^71^, which is dysfunctional in many neurodegenerative diseases, including AD. Hence, the association between tRFs of mitochondrial origin that carry motifs complementary to those of cholinergic transcripts, resemble the enriched sequence-based motifs to cholinergic transcripts in MT-tRFs from the cerebrospinal fluid (CSF) and blood of donors with Parkinson’s Disease (PD)^72^. These results suggest that the mitochondrial damage in neurodegenerative disorders such as PD and AD relates to the failed regulation of cholinergic targets by CholinotRFs, and potentially to a broader cholinergic and mitochondrial linked defect.

Surprisingly, we observed mRNA changes associated with ACh production and cholinergic regulation in the hypothalamus and the STG, although these regions lack cholinergic neurons^63^. However, previous reports presented AD-related cholinergic changes in oligodendrocytes^64^, astrocytes^65^ and microglia^14^. Therefore, we asked if the observed STG bulk RNA-Seq changes occurred in non-neuronal populations. Indeed, the MTG of AD donors compared to apparently healthy aged controls was compatible with contribution of different cell populations to the identified changes, and the linear combinations representing gene expression in the MTG were consistent with the altered STG bulk RNA-Seq.

In both AD females and males, our MTG small RNA changes were attributed to the alterations in cell numbers, rather than expression changes in each cell population. Of those, the relative percentage of affected cells was of oligodendrocytes, astrocytes and microglia-PVMs in both AD females and males. Specifically, non-neuronal populations were enlarged in cognitively impaired females compared to controls, contrasting with a reduction in cognitively impaired males. Furthermore, the cholinergic transcripts whose levels were modified in our bulk RNA-Seq could be attributed to MTG glia. Others have reported that vulnerable oligodendrocytes in AD may induce myelin breakdown and loss of the myelin sheath, which might initiate the earliest stage of AD prior to appearance of amyloid and tau pathology^64^. Astrocytes also contribute to neuroinflammation in neurodegeneration^73^. In addition, PVMs may contribute to clearance of brain neuritic plaques, and microglia play multiple AD-associated effects. Therefore, the changes of these gene targets in MTG non-neuronal populations of donors diagnosed with AD provide a potential mechanistic explanation that supports and validates those reports, as well as the changes in STG cholinergic mRNAs. Interestingly, the observed changes were more prominent in males, consistent with the observation that in AD, the cortex is affected more severely in males compared with females^67^.

Linear regression verified consistent, albeit non perfectly fit AD-related changes of cholinergic mRNAs between the two regions. This is compatible with the extraction of RNA for scRNA-Seq from the nucleus alone, missing transcripts from other areas of the cell (e.g. dendrites); also, the small RNA amounts in single cells require many rounds of amplification prior to sequencing, leading to strong amplification bias and dropouts of genes^74^; hence, scRNA-Seq data exhibit high variability between technical replicates, influencing the overall transcript levels^75^.

To challenge CholinotRFs’ roles in cholinergic regulation, we analyzed small RNA-Seq data from cell lines of female and male origins after cholinergic differentiation^51^, expecting elevated CholinotRF levels. Indeed, 4 days exposure to neurokines increased CholinotRF levels in differentiated cell lines of both female and male origins, compared to the original cell lines, supporting their role in cholinergic regulation. Intriguingly, male-originated cells showed elevated CholinotRF levels compared to non-cholinergic tRFs 2 days under neurokines exposure, but a larger effect was observed after 4 days. However, female cells showed reduced levels of most CholinotRFs after 2 days, supporting the sex-specific CholinotRF changes in the AD brains.

Cholinesterase inhibitors remain the main current strategy for treating AD symptoms. The NAc is targeted by therapeutic muscarinic agonists, which have shown some promise in clinical trials. Also, nicotinic receptor agonists modify various NAc activities. However, ChEIs may cause serious side effects, especially in females^7,16^; including exacerbated anticholinergic burden^15^ highlighting the need for more finely tuned ChEI treatments. Correspondingly, our findings point to considering cholinergic impairments in female brains upon ChEIs prescription. Furthermore, our findings revealed previously unknown correlation between tRF declines predicted to target cholinergic genes with the female- and NAc-specific elevation of those mRNA levels. Therefore, small RNA regulators of the cholinergic tone in the diseased brain. Whether such changes also occur in the nucleus basalis of Meynert remains to be determined.

We acknowledge limitations of our study, which did not define precise implications of particular small RNA changes. Expression data from cell lines of female- and male-origin before and after cholinergic differentiation only reflect one cell type and a single mechanism underlying the cholinergic changes in AD, calling for more complex model systems and revealing the need for investigation of the molecular mechanisms operating in the various cell populations, and adequately powered studies to address the pharmacological implications of our findings.

## Acknowledgments

The authors thank the study participants of the Religious Orders Study (ROS) and Rush Memory and Aging Project (MAP) and staff of the Rush Alzheimer’s Disease Center in Chicago, supported by grants from the National Institute of Health (NIH; P30AG10161, P30AG72975, R01AG15819, R01AG17917. U01AG46152, U01AG61356). The data available in the AD Knowledge Portal would not be possible without the participation of research volunteers and the contribution of data by collaborating researchers. This work was further supported by the Israel Science Foundation (ISF; 3213/19; to H. Soreq and Y. Loewenstein and 1016/18; to H. Soreq), the National Institute of Health (NIH; Aging grant 5P01AG014449-21 to E. Mufson and H Soreq), Keter Holdings (to H. Soreq), the Gatsby Charitable Foundation (to Y.L.) and the K.Stein foundation (to H.S). D. Shulman’s and N. Madrer’s fellowships were supported by the US friends of the Hebrew University of Jerusalem, the Leopold Sara and Norman Israel fund and the Sephardic Foundation on Aging. Sima Dubnov is an awardee of an Azrieli Foundation PhD fellowship. The figures in this article were created with BioRender.com.

