## Supplemental Figures for "Sex-specific declines in cholinergic-targeting tRNA fragments in the nucleus accumbens in Alzheimer’s disease"

### Supplementary Figures

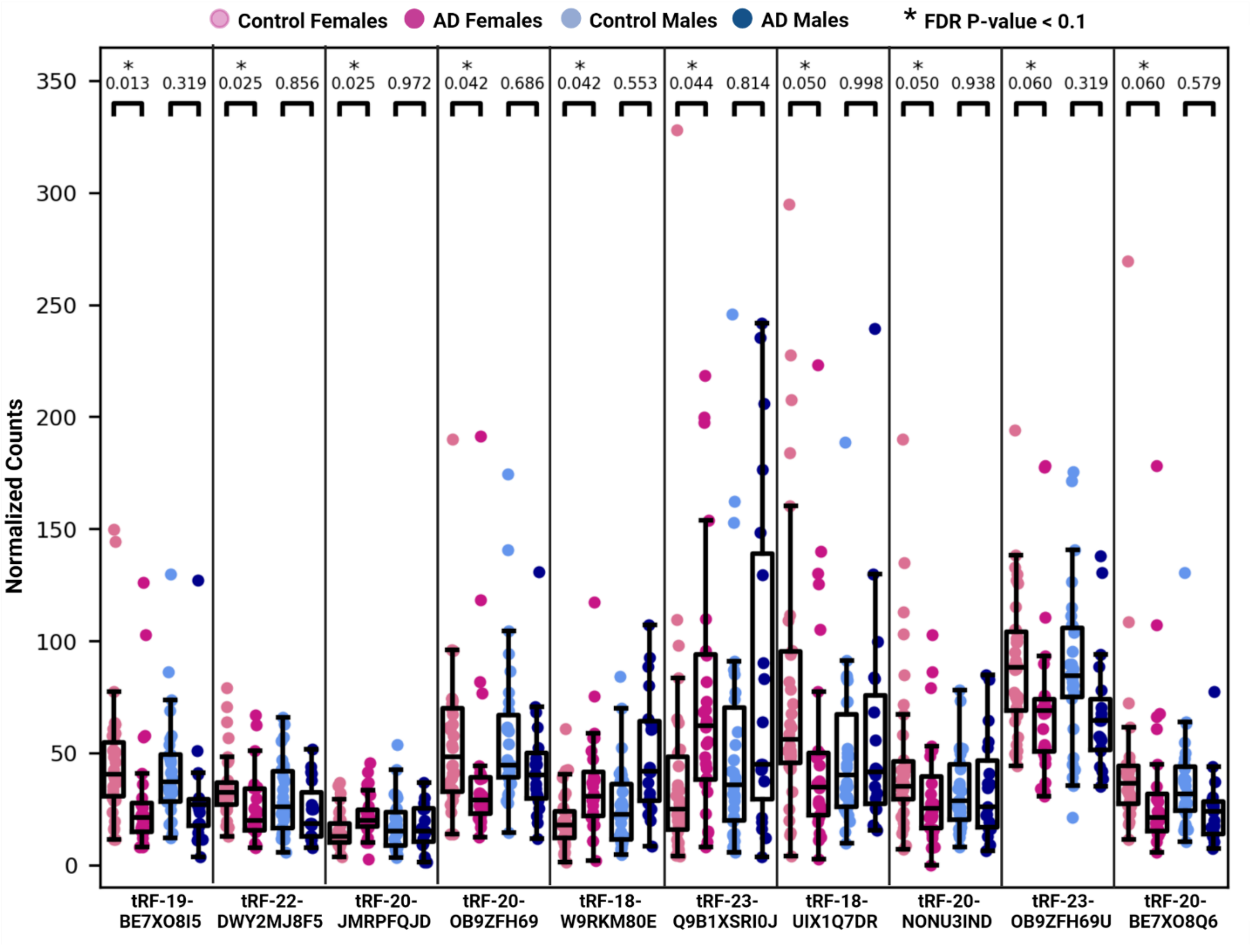


***Figure S1:*** *Boxplot plotted normalized counts of tRFs with modified levels in AD females, but not males, compared to controls. Pale pink: Female controls. Dark pink: AD females. Pale blue: Male controls. Dark blue: AD males. * changes with 0.1 > FDR corrected Kolmogorov-Smirnov P-value. Notably: 2 points with extreme values are excluded from the figure for convenient presentation: 1 AD female for tRF-23-Q9B1XSRI0J; 1 control female for tRF-18-UIX1Q7DR.*

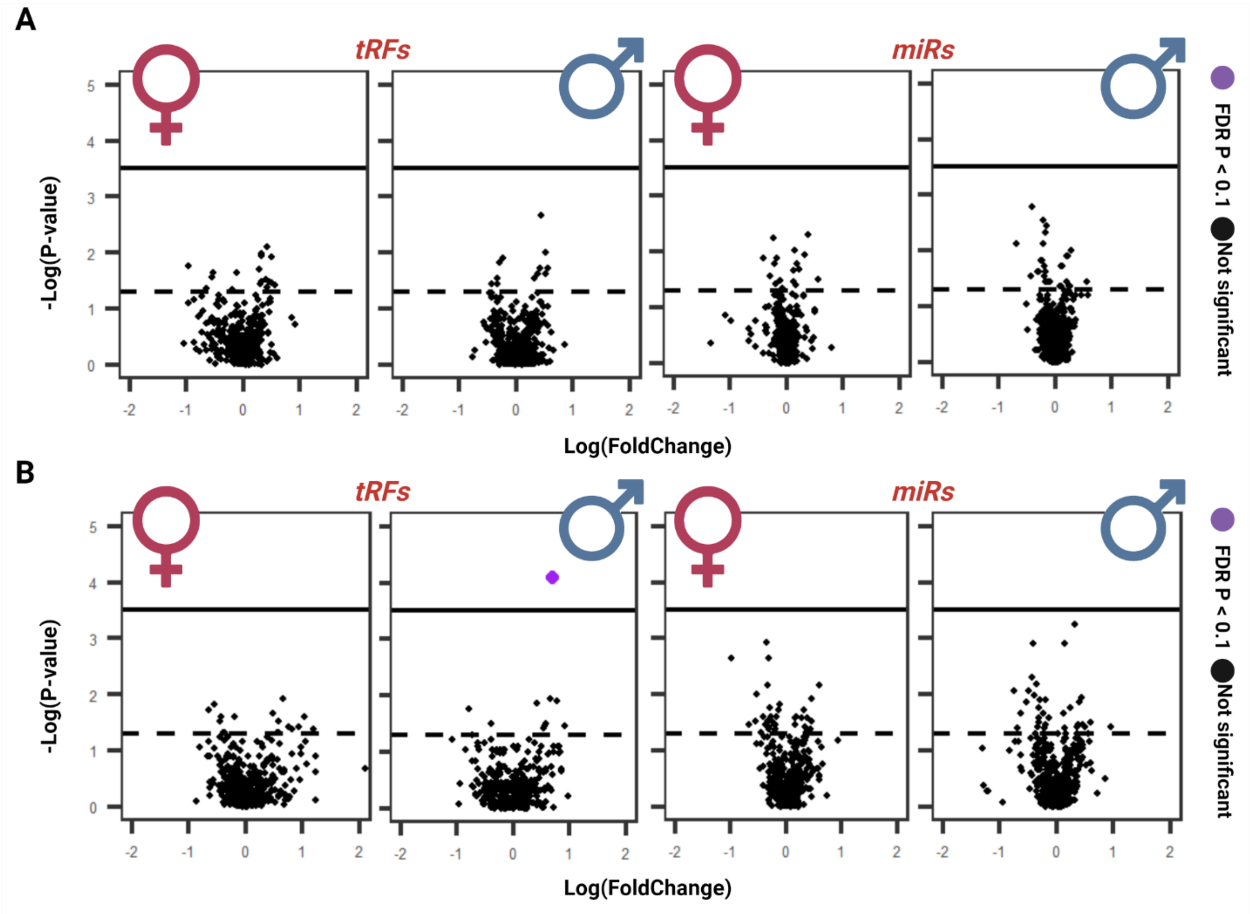


***Figure S2:*** *Kolmogorov-Smirnov P-values for NAc-expressed tRFs and miRs from female and male AD brains compared to controls by Pathophysiological measures. X axis: Log2FoldChange. Y axis: Log10(P-value). Purple points: sncRNAs with FDR-corrected P-value < 0.1. Black dashed line: p-value of 0.05. Black solid line: FDR corrected P-value < 0.1. A) Changes in samples classified by the quantity of Amyloid-β plaques. B) Changes in samples classified by the quantity of neurofibrillary tangles (NFTs).*

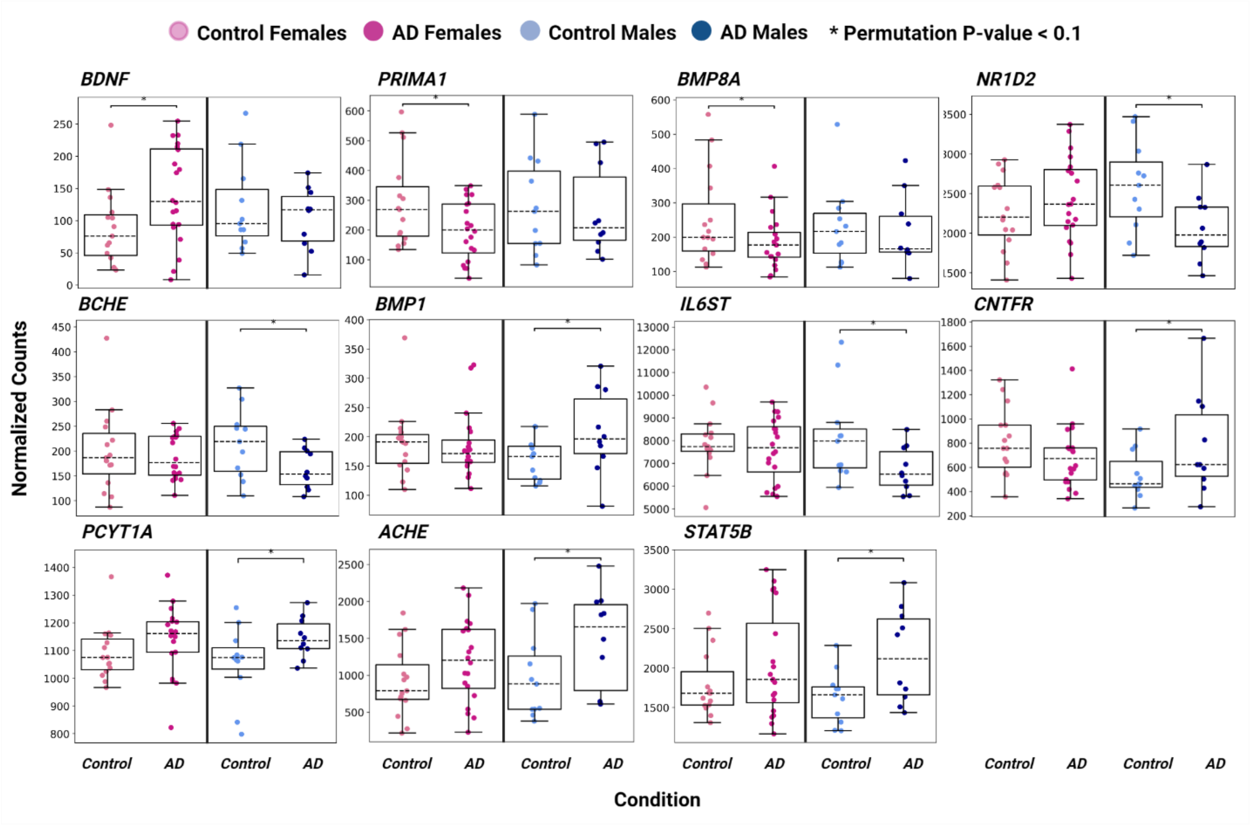


***Figure S3***: *Normalized levels of the cholinergic target mRNAs that were modified in the hypothalamus of AD compared to control groups. Pale pink: Female controls. Dark pink: AD females. Pale blue: Male controls. Dark blue: AD males. * changes with Permutation P-value < 0.1.*

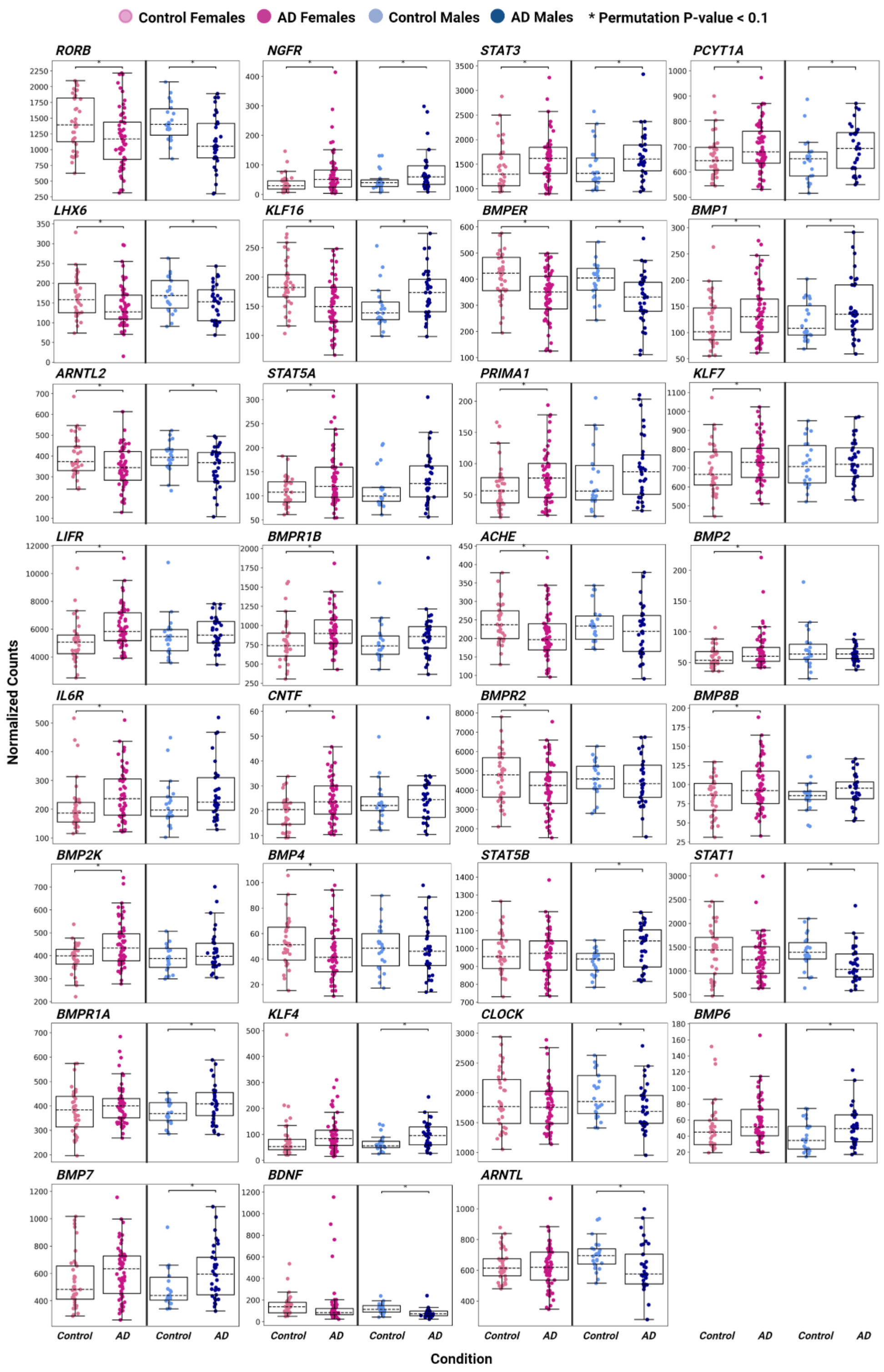


**Figure S4**: Normalized levels of the cholinergic target mRNAs that were modified in the STG of AD compared to control groups. Pale pink: Female controls. Dark pink: AD females. Pale blue: Male controls. Dark blue: AD males. * changes with Permutation P-value < 0.1.
